# Covert attention increases the gain of stimulus-evoked population codes

**DOI:** 10.1101/2020.07.30.228981

**Authors:** Joshua J. Foster, William Thyer, Janna W. Wennberg, Edward Awh

**Affiliations:** Department of Psychology, The University of Chicago, Chicago, Illinois 60637; Institute for Mind and Biology, The University of Chicago, Chicago, Illinois 60637; Department of Psychological and Brain Sciences, Boston University, Boston, Massachusetts 02215; Center for Systems Neuroscience, Boston University, Boston, Massachusetts, 02215; Department of Psychology, University of California, San Diego, La Jolla, California, 92092

## Abstract

Covert spatial attention has a variety of effects on the responses of individual neurons. However, relatively little is known about the net effect of these changes on sensory population codes, even though perception ultimately depends on population activity. Here, we measured the electroencephalogram (EEG) in human observers (male and female), and isolated stimulus-evoked activity that was phase-locked to the onset of attended and ignored visual stimuli. Using an encoding model, we reconstructed spatially selective population tuning functions from the pattern of stimulus-evoked activity across the scalp. Our EEG-based approach allowed us to measure very early visually evoked responses occurring ~100 ms after stimulus onset. In Experiment 1, we found that covert attention increased the amplitude of spatially tuned population responses at this early stage of sensory processing. In Experiment 2, we parametrically varied stimulus contrast to test how this effect scaled with stimulus contrast. We found that the effect of attention on the amplitude of spatially tuned responses increased with stimulus contrast, and was well-described by an increase in response gain (i.e., a multiplicative scaling of the population response). Together, our results show that attention increases the gain of spatial population codes during the first wave of visual processing.

**Significance Statement:** We know relatively little about how attention improves population codes, even though perception is thought to critically depend on population activity. In this study, we used an encoding-model approach to test how attention modulates the spatial tuning of stimulus-evoked population responses measured with EEG. We found that attention multiplicatively scales the amplitude of spatially tuned population responses. Furthermore, this effect was present within 100 ms of stimulus onset. Thus, our results show that attention improves spatial population codes by increasing their gain at this early stage of processing.

## Introduction

Covert spatial attention improves perception by improving neural representations in visual cortex (Maunsell, 2015; Sprague et al., 2015). At the level of individual neurons, spatial attention not only increases the amplitude of responses (Luck et al., 1997; McAdams and Maunsell, 1999), but also has a variety of effects on the spatial tuning of neurons: receptive fields shift toward attended locations, and attention increases the size of the receptive field of some neurons while decreasing the size of others (Connor et al., 1997; Womelsdorf et al., 2006, 2008; Anton-Erxleben et al., 2009; for reviews, see Anton-Erxleben and Carrasco, 2013; Sprague et al., 2015). Ultimately, however, perception depends on the joint activity of large ensembles of cells (Pouget et al., 2000). Thus, there is strong motivation to understand the net effect of these local changes for population representations (Sprague et al., 2015).

There is clear evidence that attended stimuli evoke larger population responses than unattended stimuli. For instance, covert attention increases the amplitude of visually evoked potentials measured with electroencephalography (EEG; e.g. van Voorhis and Hillyard, 1977; Itthipuripat et al., 2014a), which reflect the aggregate activity of many neurons (Nunez and Srinivasan, 2006). However, studies that measure changes in the overall amplitude of population responses do not reveal how attention influences the *information content* of population activity (Serences and Saproo, 2012). Thus, researchers have turned to multivariate methods. Sprague and Serences (2013), for example, used an inverted encoding model (IEM) to reconstruct population-level representations of stimulus position from patterns of activity measured with functional magnetic resonance imaging (fMRI). They found that spatially attending a stimulus increased the amplitude of spatial representations across the visual hierarchy without reliably changing their size (also see Vo et al., 2017; Itthipuripat et al., 2019; but see Fischer and Whitney, 2009).

Although fMRI is a powerful tool for assaying population codes, two major limitations prevent clear conclusions regarding the effect of attention on stimulus-driven activity. First, the sluggish blood-oxygen-level-dependent (BOLD) signal that is measured with fMRI provides little information about *when* attention modulates population codes. Second, growing evidence suggests that the effect of attention on the BOLD signal does not reflect a modulation of the stimulus-evoked response at all, but instead reflects a stimulus-independent shift in baseline activity. These studies varied stimulus contrast to measure neural contrast-response functions (CRFs), which can be modulated by attention in several ways (Fig. 1). Whereas unit-recording and EEG studies have found that attentional modulation of neural responses depends on stimulus contrast, either multiplicatively scaling the CRF (*response gain*, Fig. 1a) or shifting the CRF to the left (*contrast gain*, Fig. 1b) (Reynolds et al., 2000; Martinez-Trujillo and Treue, 2002; Kim et al., 2007; Itthipuripat et al., 2014a, 2014b, 2019), fMRI studies have found that spatial attention increases the BOLD signal in visual cortex by the same amount regardless of stimulus contrast, even when no stimulus is presented at all (an *additive shift*, Fig. 1c; Buracas and Boynton, 2007; Murray, 2008; Pestilli et al., 2011; Sprague et al., 2018b; Itthipuripat et al., 2019; but see Li et al., 2008). This finding suggests that the effect of attention on the BOLD response reflects top-down inputs to visual cortex rather than a modulation of stimulus-driven activity (Murray, 2008; Itthipuripat et al., 2014a). Therefore, extant work has not yet determined how attention changes *stimulus-driven* population codes.

**Figure 1.**
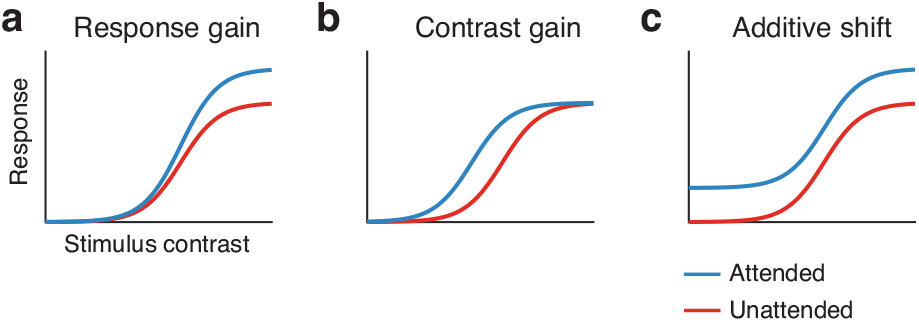
Attentional modulations of contrast-response functions (CRFs). Each plot shows the level of the sensory activity as a function of stimulus contrast and attention. Three kinds of attentional modulation have been reported in past studies. **(a)** Response gain: attention multiplicatively scales the CRF, such that attention has a larger effect at higher stimulus contrasts. **(b)** Contrast gain: attention shifts the CRF to the left, increasing the effective strength of the stimulus. **(c)** Additive shift: attention shifts the entire CRF up. Because an additive shift increases neural activity in the absence of a visual stimulus (i.e. stimulus contrast of 0%), additive shifts likely reflects a top-down attention-related signal rather than a modulation of stimulus-driven activity.

Here, we used EEG to examine how spatial attention modulates the spatial tuning of stimulus-driven population responses. We measured *stimulus-evoked* activity (i.e., activity that is phase-locked to stimulus onset) to isolate the stimulus-driven response from ongoing activity that is independent of the stimulus. We used an IEM (Brouwer and Heeger, 2009) to reconstruct spatially selective channel-tuning functions (CTFs) from the pattern of stimulus-evoked activity across the scalp. The resulting CTFs reflect the spatial tuning of the population activity that is measured with EEG. We focused our analysis in an early window, approximately 100 ms after stimulus onset. Activity at this latency is thought to primarily reflect the first wave of sensory activity evoked by a stimulus in extrastriate cortex (Clark and Hillyard, 1996; Martínez et al., 1999). In Experiment 1, we found that attention increased the amplitude of stimulus-evoked CTFs. Thus, attention increased the gain of spatial population codes at this early stage of sensory processing. In Experiment 2, we further characterized the effect of attention on spatial population codes by parametrically varying stimulus contrast. We found that the effect of attention on the amplitude of stimulus-evoked CTFs increased with stimulus contrast, and was well-described as an increase in response gain (Fig. 1a). Taken together, our results show that attention increases the gain of stimulus-evoked population codes at early stages of sensory processing.

## Materials and Methods

### Subjects

Forty-five volunteers (21 in Experiment 1and 24 in Experiment 2) participated in the experiments for monetary compensation ($15/hr). Subjects were between 18 and 35 years old, reported normal or corrected-to-normal visual acuity, and provided informed consent according to procedures approved by the University of Chicago Institutional Review Board.

#### Experiment 1

Our target sample size was 16 subjects in Experiment 1, following our past work using an IEM to reconstruct spatial CTFs from EEG activity (Foster et al., 2016). Twenty-one volunteers participated in Experiment 1 (8 male, 13 female; mean age = 22.7 years, SD = 3.2). Four subjects were excluded from the final sample for the following reasons: we were unable to prepare the subject for EEG (n = 1); we were unable to obtain eye tracking data (n = 1); the subject did not complete enough blocks of the task (n = 1); and residual bias in eye position (see Eye movement controls) was too large (n = 1). The final sample size was 17 (6 male, 11 female; mean age = 22.7 years, SD = 3.4). We overshot our target sample size of 16 because the final subject was scheduled to participate before we reached our target sample size.

#### Experiment 2

In Experiment 2, we increased our target sample size to 20 subjects to increase statistical power because we sought to test how the effect of attention changes with stimulus contrast. Twenty-four volunteers participated in Experiment 2 (6 male, 16 female; mean age = 24.0 years, *SD* = 3.0), four of which had previously participated in Experiment 1. For four subjects, we terminated data collection and excluded the subject from the final sample for the following reasons: we were unable to obtain eye tracking data (*n* = 1); the subject had difficulty performing the task (*n* = 1); the subject made too many eye movements (*n* = 2). The final sample size was 20 (5 male, 15 female; mean age = 24.0 years, *SD* = 2.8).

### Apparatus and stimuli

We tested the subjects in a dimly lit, electrically shielded chamber. Stimuli were generated using Matlab (MathWorks, Natick, MA) and the Psychophysics Toolbox (Brainard, 1997; Pelli, 1997). Subjects viewed the stimuli on a gamma-corrected 24” LCD monitor (refresh rate: 120 Hz, resolution 1080 x 1920 pixels) with their chin on a padded chin rest (viewing distance: 76 cm in Experiment 1, 75 cm in Experiment 2). Stimuli were presented against a mid-gray background (~61 cd/m^2^).

### Task procedures

On each trial, observers viewed a sequence of four bullseye stimuli (Fig. 2a). Across blocks, we manipulated whether observers attended the bullseye stimuli (*attend-stimulus* condition) or attended the central fixation dot (*attend-fixation* condition). In the attend-stimulus condition, observers monitored the sequence for one bullseye that was lower contrast than the rest (a *bullseye target).* In the attend-fixation condition, observers monitored the fixation dot for a 100-ms decrement in contrast (a *fixation target).* Contrast decrements for both the bullseye targets and fixation targets occurred on half of the trials in both conditions, and the trials that contained bullseye targets and fixation targets were determined independently. We instructed subjects to disregard changes in the unattended stimulus. Although past work has suggested that there may be differences in the cortical regions that support attention to peripheral locations and attention to fixated locations (Kelley et al., 2008), we contrasted target-evoked responses in these conditions because of the powerful effect that this manipulation of attention has on stimulus-evoked responses. Furthermore, recent studies that have used fMRI to examine the effect of attention on spatially tuned population responses have manipulated attention in the same way (e.g. Sprague and Serences, 2013; Itthipuripat et al., 2019). Therefore, this manipulation of attention allows for comparison with these past studies.

**Figure 2.**
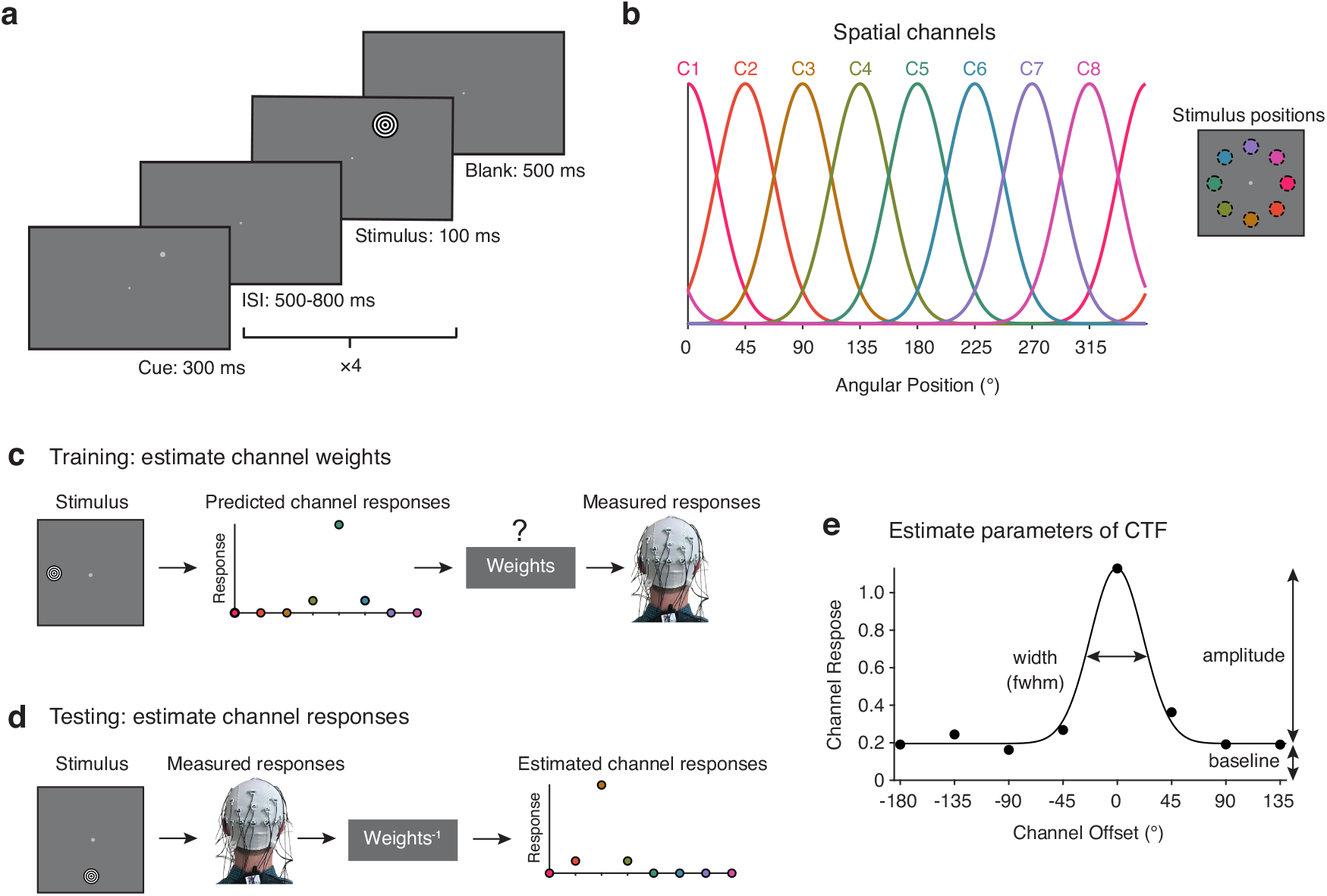
Experimental task and inverted encoding model method. **(a)** Human observers viewed a series of four bullseye stimuli, each separated by a variable inter-stimulus interval (ISI). The trial began with a peripheral cue that indicated where the bullseye stimuli would appear. In the attend-stimulus condition, observers monitored the bullseye stimuli for one stimulus that was lower contrast than the others. In the attend-fixation condition, observers - monitored the fixation dot for a brief reduction in contrast. **(b)** We modelled power at each electrode as the weighted sum of eight spatially selective channels (here labeled C1-C8). Each channel was tuned for one of the eight positions at which the stimuli could appear in the experiment (shown on the right). The curves show the predicted response of the eight channels as a function of stimulus position (i.e. the basis set). **(c)** In the training phase of the analysis, the predicted channel responses (determined by the basis set) served as regressors, allowing us to estimate a set of channel weights that specified the contribution of each spatial channel to power measured at each electrode. **(d)** In the testing phase of the analysis, we used the channel weights from the training phase to estimate the response of each channel given an independent test set of data. **(e)** We circularly shifted the channel response profiles for each stimulus position to a common center and averaged them to obtained a channel tuning function (CTF) shown as black circles (data simulated for illustrative purposes). A Channel Offset of 0° on the x-axis marks the channel tuned for the location of the stimulus. We fitted an exponentiated cosine function to CTFs to measure their *amplitude, baseline*, and *width* (measured as full-width-at-half-maximum or fwhm).

Observers fixated a central dot (0.1° in diameter, 56.3% Weber contrast, i.e. 100 × (L - L_b_)/L_b_, where L is stimulus luminance and L_b_ is the background luminance) before pressing spacebar to initiate each trial. Each trial began with a 400 ms fixation display. A peripheral cue (0.25° in diameter, 32.8% Weber contrast) was presented where the bullseye stimuli would appear for 300 ms. On each trial, the bullseyes appeared at one of eight locations equally spaced around fixation at an eccentricity of 4°. Each bullseye (1.6° in diameter, 0.12 cycles/°) appeared for 100 ms. The cue and each of the bullseyes were separated by a variable inter-stimulus interval between 500 and 800 ms. Bullseye targets (the bullseye that was lower contrast than the others) were never the first bullseye in the sequence. Thus, the first bullseye of each trial established the pedestal contrast the trial (i.e., the contrast of the non-target bullseyes). Fixation targets (a 100-ms decrement in the contrast of the fixation dot) occurred at the same time as one of the bullseye stimuli, and like bullseye targets, fixation targets never occurred during the presentation of the first bullseye of the trial. Both bullseye and fixation targets occurred on 50% of trials, determined randomly and independently for each stimulus to preclude accurate performance based on attention to the wrong aspect of the display. On trials with both a bullseye target and fixation target (25% of trials), the timing of each target was determined independently, such that the targets co-occurred on approximately 33% of these trials. The final bullseye in each trial was followed by a 500 ms blank display before the response screen appeared. Each trial ended with a response screen that prompted subjects to report whether or not a target was presented in the relevant stimulus. Subjects responded using the numberpad of a standard keyboard (“1” = change, “2” = no change). The subject’s response appeared above the fixation dot, and they could correct their response if they pressed the wrong key. Finally, subjects confirmed their response by pressing the spacebar.

#### Experiment 1

In Experiment 1, the pedestal contrast of the bullseye was always 89.1% Michelson contrast (100 × (L_max_ − L_min_)/(L_max_ + L_min_), where Lmax in the maximum luminance and L_min_ is the minimum luminance). Subjects completed a 3.5-hour session. The session began with a staircase procedure to adjust task difficulty (see Staircase Procedures). Subjects then completed 12-20 blocks (40 trials each) during which we recorded EEG. Thus, subjects completed between 480 and 800 trials (1920-3200 stimulus presentations). The blocks alternated between the attend-stimulus and attend-fixation conditions, and we counterbalanced task order across subjects.

#### Experiment 2

In Experiment 2, we manipulated the contrast of the bullseye stimuli. We included 5 pedestal contrasts (6.25, 12.5, 25.0, 50.0, and 90.6% Michelson contrast). Thus, there were 10 conditions in total (2 attention conditions × 5 pedestal contrasts). Subjects completed three sessions: a 2.5-hour behavior session to adjust task difficulty in each condition (see Staircase Procedures), followed by two 3.5-hour EEG sessions. All sessions were completed within a 10-day period. Each block consisted of 104 trials: eight trials for each of the 10 conditions, and an additional 12 trials in each condition at the highest pedestal contrast (90.6% contrast) for the purpose of training the encoding model (see Training and testing data). Each block included a break at the halfway point. As in Experiment 1, the blocks alternated between the attend-stimulus and attend-fixation conditions, and we counterbalanced task order across subjects. We aimed to have each subject complete 20 blocks across the EEG sessions to obtain 160 testing trials for each condition (640 stimulus presentations), and 480 training trials (1920 stimulus presentations). All subjects completed 20 blocks with the following exceptions: three subjects completed 18 blocks, and one subject completed 24 blocks.

In Experiment 2, we made one minor change from Experiment 1: the experimenter could manually provide feedback to the observer to indicate whether they noticed blinks or eye movements during the trial by pressing a key outside the recording chamber. When feedback was provided, the text “blink” or “eye movement” was presented in red for 500 ms after the observer had made their response.

### Staircase procedures

In each experiment, we used a staircase procedure to match difficulty across conditions in both experiments. We adjusted difficulty by adjusting the size of the contrast decrement for each condition independently.

#### Experiment 1

In Experiment 1, subjects completed six staircase blocks of 40 trials (three blocks for each condition) before we started the EEG blocks of the task. Thus, subjects completed 120 staircase trials for each condition. We used a 3-down-1-up procedure to adjust task difficulty: after three correct responses in a row, we reduced the size of the contrast decrement by 2%; after an incorrect response, we increased the size of the contrast decrement by 2%. This procedure was designed to hold accuracy at ~80% correct (García-Pérez, 1998). The final size of the contrast decrements in the staircase blocks were used for the EEG blocks. During the EEG blocks, we examined accuracy in each condition every four blocks (two blocks of each condition), and adjusted the size of the contrast decrements to hold accuracy as close to 80% as possible.

#### Experiment 2

In Experiment 2, subjects completed a 2.5-hour staircase session prior to the EEG sessions. We adjusted difficulty for each of the 10 conditions independently (2 attention conditions × 5 pedestal contrast). Subjects completed 16 blocks of 40 trials, alternating between the attend-fixation and attend-stimulus conditions. The five contrast levels were randomized within each block. Thus, observers completed 64 staircase trials for each of the 10 conditions. We used a weighted up/down procedure to adjust task difficulty: after a correct response, we reduced the size of the contrast decrement by 5%; after an incorrect response, we increased the size of the contrast decrement by 17.6%. This procedure held accuracy fixed at ~76%. The staircase procedure continued to operate throughout the EEG sessions.

### EEG acquisition

We recorded EEG activity from 30 active Ag/AgCl electrodes mounted in an elastic cap (Brain Products actiCHamp, Munich, Germany). We recorded from International 10-20 sites: Fp1, Fp2, F7, F3, Fz, F4, F8, FT9, FC5, FC1, FC2, FC6, FT10, T7, C3, Cz, C4, T8, CP5, CP1, CP2, CP6, P7, P3, Pz, P4, P8, O1, Oz, O2. Two additional electrodes were affixed with stickers to the left and right mastoids, and a ground electrode was placed in the elastic cap at position Fpz. All sites were recorded with a right-mastoid reference and were re-referenced offline to the algebraic average of the left and right mastoids. We recorded electrooculogram (EOG) data using passive electrodes, with a ground electrode placed on the left cheek. Horizontal EOG was recorded from a bipolar pair of electrodes placed ~1 cm from the external canthus of each eye. Vertical EOG was recorded from a bipolar pair of electrodes placed above and below the right eye. Data were filtered online (low cut-off = .01 Hz, high cut-off = 80 Hz, slope from low-to high-cutoff = 12 dB/octave), and were digitized at 500 Hz using BrainVision Recorder (Brain Products, Munich, German) running on a PC. Impedance values were kept below 10 kΩ.

### Eye tracking

We monitored gaze position using a desk-mounted EyeLink 1000 Plus infrared eye-tracking camera (SR Research, Ontario, Canada). Gaze position was sampled at 1000 Hz. Head position was stabilized with a chin rest. According to the manufacturer, this system provides spatial resolution of 0.01° of visual angle, and average accuracy of 0.25-0.50° of visual angle. We calibrated the eye tracker every 1 −2 blocks of the task, and between trials during the blocks if necessary. We drift-corrected the eye tracking data for each trial by subtracting the mean gaze position measured during a 200 ms window immediately before the onset of the cue.

### Artifact rejection

We excluded data from some electrodes for some subjects because of low quality data (excessive high-frequency noise or sudden steps in voltage). In Experiment 1, we excluded one or two electrodes for three subjects in our final sample. In Experiment 2, we excluded electrodes Fp1 and Fp2 for all subjects because we obtained poor-quality data (high-frequency noise and slow drifts) at these sites for most subjects, and we excluded data for one additional electrode for two subjects in our final sample. In both experiments, all excluded electrodes were located at frontal or central sites. Our window of interest was from 200 ms before stimulus onset until 500 ms after stimulus onset. We segmented the EEG data into epochs time-locked to the onset of each bullseye stimulus (starting 1200 ms before stimulus onset and ending 1500 ms after stimulus onset). We segmented data into longer epochs so that the epochs were long enough to apply a high-pass filter (see Evoked power), and so that our window of interest was not contaminated with edge artifacts when filtering the data. We baselined corrected the EEG data by subtracting mean voltage during the 200-ms window immediately prior to stimulus onset. We visually inspected the segmented EEG data for artifacts (amplifier saturation, excessive muscle noise, and skin potentials), and the eye tracking data for ocular artifacts (blinks, eye movements, and deviations in eye position from fixation), and discarded any epochs contaminated by artifacts. In Experiment 1, all subjects included in the final sample had at least 800 artifact-free epochs for each condition. In Experiment 2, all subjects included in the final sample had at least 450 artifact-epochs for testing the IEM in each condition, and at least 1500 artifact-free epochs for training the IEM (see Training and Test Data).

### Eye movement controls

After artifact rejection, for each subject we inspected mean gaze position as a function of stimulus position for the attend-stimulus and attend-fixation conditions separately. For all subjects in the final samples, mean gaze position varied by less than 0.2° of visual angle across stimulus positions. One subject in Experiment 1 was excluded from the final sample because they did not meet this criterion. To verify that removal of ocular artifacts was effective, we inspected mean gaze position (during the 100-ms presentation of each stimulus) as a function of stimulus position for the attend-stimulus and attend-fixation conditions separately. In both experiments, we observed very little variation in mean gaze position (across subjects) as a function of stimulus position (< 0.05° of visual angle) for both the attend-stimulus and attend-fixation conditions (Figure 3), confirming that we achieved an extremely high standard of fixation compliance after epochs with artifacts were removed. Thus, the effects of attention reported below cannot be attributed to variation in eye position.

**Figure 3.**
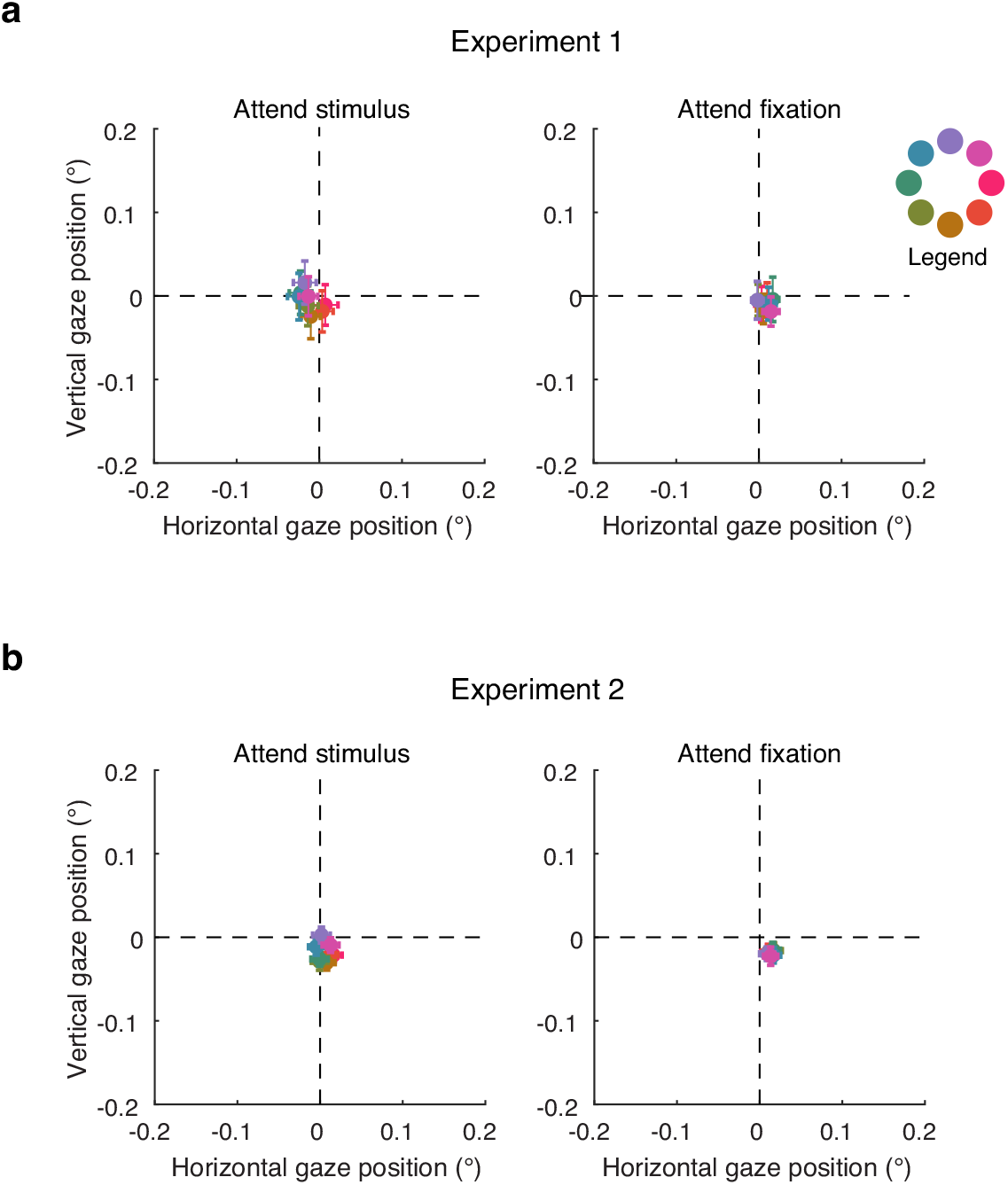
Residual variation in eye position after artifact rejection. **(a)** Mean gaze coordinates in Experiment 1 as a function of stimulus position for the attend-stimulus (left) and attend-fixation (right) conditions. Gaze coordinates were calculated during the 100-ms presentations of the bullseye stimuli (averaging across the four presentations in the trial sequence). **(b)** Same for Experiment 2. The legend at the right of the plot shows which color corresponds to each of the eight stimulus positions. Error bars show ±1 SEM across subjects.

### Controlling for stimulus contrast

On half of trials, one of the four bullseyes was lower contrast than the rest (i.e. a target). Thus, the average contrast of the bullseyes was slightly lower than the pedestal contrast (i.e. the contrast of the non-target bullseyes), and small differences in average contrast may have emerged between conditions after rejection of data that were contaminated by EEG artifacts or eye movements. However, the difference in mean contrast between the attend-stimulus and attend-fixation conditions after artifact rejection was negligible. In Experiment 1, mean contrast of the bullseye stimuli was 87.4% (*SD* = 0.97) in the attend-stimulus condition and 87.5% (*SD* = 0.92) in the attend-fixation condition. Similarly, in Experiment 2, the mean contrast of the bullseye stimuli was comparable for the attend-stimulus and attend-fixation conditions for all pedestal contrasts (Table 1).

**Table 1.**
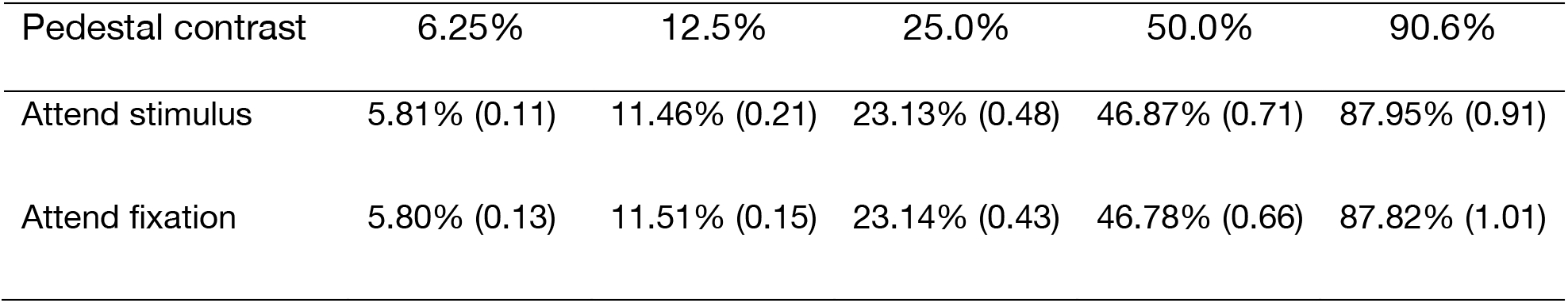
Mean Michelson contrast (and standard deviation) of the bullseye in Experiment 2 as a function of task condition and pedestal contrast of the bullseye stimuli.

### Evoked power

A Hilbert Transform (Matlab Signal Processing Toolbox) was applied to the segmented EEG data to obtain the complex analytic signal, *z(t)*, of the EEG, *f(t)*:

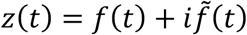

where 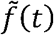 is the Hilbert Transform of *f(t)*, and 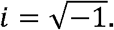. The complex analytic signal

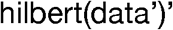

was extracted for each electrode using the following Matlab syntax: where data is a 2D matrix of segmented EEG (number of trials × number of samples). We calculated *evoked* power by first averaging the complex analytic signals across trials, and then squaring the complex magnitude of the averaged analytic signal. Evoked power isolates activity phase-locked to stimulus onset because only activity with consistent phase across trials remains after averaging the complex analytic signal across trials. Trial averaging was performed for each stimulus position separately within each block of training or test data for the IEM analyses (see Training and testing data).

For some analyses, we high-pass filtered the data with a low-cutoff of 4-Hz to remove low frequency activity before calculating evoked power. We used EEGLAB’s “eegfilt.m” function (Delorme and Makieg, 2004), which implements a two-way leastsquares finite impulse response filter. This filtering method uses a zero-phase forward and reverse operation, which ensures that phase values are not distorted, as can occur with forward-only filtering methods.

### Alpha-band power

To calculate alpha-band power at each electrode, we bandpass filtered the raw EEG data between 8 and 12 Hz using the “eegfilt.m” function in EEGLAB (Delorme and Makieg, 2004), and applied a Hilbert transform (MATLAB Signal Processing Toolbox) to the bandpass-filtered data to obtain the complex analytic signal. Instantaneous power was calculated by squaring the complex magnitude of the complex analytic signal.

### Inverted encoding model

We used an inverted encoding model (Brouwer and Heeger, 2009, 2011) to reconstruct spatially selective channel-tuning functions (CTFs) from the distribution of power across electrodes (Foster et al., 2016). We assumed that the power at each electrode reflects the weighted sum of eight spatially selective channels (i.e., neuronal populations), each tuned for a different angular position (Fig. 2b). We modeled the response profile of each spatial channel across angular locations as a half sinusoid raised to the twenty-fifth power:

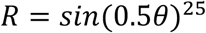

where θ is angular location (0–359°), and *R* is the response of the spatial channel in arbitrary units. This response profile was circularly shifted for each channel such that the peak response of each spatial channel was centered over one of the eight locations at which the bullseye stimuli could appear (0°, 45°, 90°, etc.).

An IEM routine was applied to each time point. We partitioned our data into independent sets of training data and test data (see Training and testing data). The analysis proceeded in two stages (training and test). In the training stage (Fig. 2c), training data (*B_1_)* were used to estimate weights that approximate the relative contribution of the eight spatial channels to the observed response measured at each electrode. Let *B*_1_ (*m* electrodes × *n_1_* measurements) be the power at each electrode for each measurement in the training set, *C*_1_ (*k* channels × *n*_1_ measurements) be the predicted response of each spatial channel (determined by the basis functions, see Fig. 2b) for each measurement, and *W* (*m* electrodes × *k* channels) be a weight matrix that characterizes a linear mapping from “channel space” to “electrode space”. The relationship between *B*_1_, *C*_1_, and *W* can be described by a general linear model of the form:

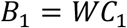

The weight matrix was obtained via least-squares estimation as follows:

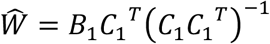

In the test stage (Fig. 2d), we inverted the model to transform the observed test data *B*_2_ (*m* electrodes × *n_2_* measurements) into estimated channel responses, *C*_2_ (*k* channels × *n_2_* measurements), using the estimated weight matrix, *Ŵ*, that we obtained in the training phase:

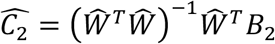

Each estimated channel response function was then circularly shifted to a common center, so the center channel was the channel tuned for the position of the probed stimulus (i.e., 0° on the “Channel Offset” axes), then averaged these shifted channel-response functions across the eight stimulus locations to obtain a CTF. Finally, because the exact contributions of each spatial channel to each electrode (i.e., the channel weights, *W*) likely vary across subjects, we applied the IEM routine separately for each subject.

### Training and testing data

For the IEM analysis, we partitioned artifact-free epochs into three independent sets: two training sets and one test set. Within each set, we calculated power across the epochs for each stimulus position to obtain a matrix of power values across all electrodes for each stimulus position (electrodes × stimulus positions, for each time point). We equated the number of epochs for each stimulus position in each set. Some excess epochs were not assigned to any set because of this constraint. Thus, we used an iterative approach to make use of all available epochs. For each of 500 iterations, we randomly partitioned the data into training and test data (see below for details of how data partitioned into training and test sets in each experiment), and we averaged the resulting CTFs across iterations.

#### Experiment 1

When comparing CTF parameters across conditions, it is critical to estimate a fixed encoding model (i.e., train the encoding model on a common training set) that is then used to reconstruct CTFs for each condition separately (for discussion of this issue, see Sprague et al., 2018a, 2019). Thus, for Experiment 1, we estimated the encoding model using a training set that included equal numbers of trials from each condition. Note that while we trained our encoding model on a mix of the attend-stimulus and attend-fixation conditions, training on a mix of data from both conditions is not necessary for the purposes of estimating the encoding model. Rather, what is critical is to estimate channel weights just once using the same training set, so that the reconstructed CTFs for each condition can be compared on an equal footing (Sprague et al., 2018a, 2019). We opted to use a mix of the two conditions for estimated the encoding model so that observers were not completing considerably more trials in one attention condition than in the other. Specifically, in Experiment 1 we partitioned data for each condition (attend-stimulus and attend-fixation) into three sets (with the constraint that the number of trials per location in each set was also equated across conditions). We obtained training data by combining data across the two conditions before calculating power, resulting in two training sets that included equal numbers of trials from each condition. We then tested the model using the remaining set of data for each condition separately. Thus, we used the same training data to estimate a single encoding model, and varied only the test data that was used to reconstruct CTFs for each condition.

#### Experiment 2

In Experiment 2, we included additional trials in the 90.6% contrast conditions (half from the attend-stimulus condition and half from the attend-fixation condition) to train the encoding model (see Task Procedures, Experiment 2). We used high-contrast stimuli to estimate channel weights because high-contrast stimuli should drive a strong stimulus-evoked response. For each iteration of the analysis, we partitioned this data into two training sets, and generated a single testing set for each of the 10 conditions separately. We equated the number of trials included for each stimulus position in each of the testing sets.

### Quantifying changes in channel-tuning functions

To characterize how CTFs changes across conditions, we fitted CTFs with an exponentiated cosine function (Fig. 2e) of the form:

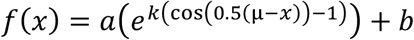

where *x* is channel offset (−180°, −135°, −90° ..., 135°). We fixed the *μ* parameter, which determines the center of the tuning function, at a channel offset of 0° such that the peak of the function was fixed at the channel tuned for the stimulus position). The function had three free parameters: *baseline (b)*, which determines the vertical offset of the function from zero; *amplitude* (*a*), which determines the height of the peak of the function above baseline; and, *concentration* (*k*) which determines the width of the function. We fitted the function with a general linear model combined with a grid search procedure (Ester et al., 2015). We converted report the concentration as width measured as full-width-at-half-maximum (fwhm): the width of the function in angular degrees halfway between baseline and the peak.

We used a subject-level resampling procedure to test for differences in the parameters of the fitted function across conditions. We drew 100,000 bootstrap samples, each containing *N*-many subjects sampled with replacement, where *N* is the sample size. For each bootstrap sample, we fitted the exponentiated cosine function described above to the mean CTF across subjects in the bootstrap sample.

In Experiment 1, to test for differences between conditions in each parameter, we calculated the difference for the parameter between the attend-stimulus and attend-fixation conditions for each bootstrap sample, which yielded a distribution of 100,000 values. We tested whether these difference distributions significantly differed from zero in either direction, by calculating the proportion of values > or < 0. We doubled the smaller value to obtain a 2-sided *p* value.

In Experiment 2, for each parameter we tested for main effects of attention and contrast, and for an attention × contrast interaction. To test for a main effect of attention, we averaged parameter estimates across contrast levels for each bootstrap sample, and calculated the difference in each parameter estimate between attention conditions for each bootstrap sample. We tested whether these difference distributions significantly differed from zero in either direction, by calculating the proportion of values > or < 0. To test for a main effect of contrast, we averaged the parameter estimates across the attention conditions, and fitted a linear function to the parameter estimates as a function of contrast. For each bootstrap sample, we calculated the slope of the best-fit linear function. We tested whether the resulting distribution of slope values significantly differed from zero in either direction by calculating the proportion of values > or < 0. Finally, to test for an attention × contrast interaction, we fitted a linear function to the parameter estimates as a function of contrast for the attend-stimulus and attend-fixation conditions separately. For each bootstrap sample, we calculated the difference in the slope of these functions between the attend-stimulus and attend-fixation conditions. We tested whether the resulting distribution of differences-in-slope values significantly differenced from zero differed from zero in either direction by calculating the proportion of values > or < 0. For both main effects and the interaction, we doubled the smaller *p* value to obtained a 2-sided *p* value.

### Quantifying contrast-response functions

We found that the effect of attention of the amplitude of stimulus-evoked CTFs varied with stimulus contrast. To further characterize this effect, we fitted the amplitude of stimulus-evoked CTFs across stimulus contrasts for each condition with a Naka-Rushton of the form:

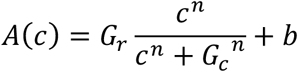

where *A* is the amplitude of stimulus-evoked CTFs, and *c* is stimulus contrast. The function had four free parameters: baseline (*b*), which determines the offset of the function from zero, response gain (*G_r_*), which determines how far the function rises above baseline, contrast gain (*G_c_*), which determines the semi-saturation point, and an exponent (n) that determines the slope of the function. We used Matlab’s “fmincon” function to minimize the sum of squared errors between the data and the Naka-Rushton function. We restricted the *b* and *G_r_* parameters to be between 0 and 10 (with 10 being a value that far exceeds the observed amplitudes of stimulus-evoked CTFs), *G_c_* to be between 0 and 100% contrast, and *n* to be between 0.1 and 10. As Itthipuripat et al. (2019) have pointed out, in the absence of a saturating function, one might obtain unrealistically estimates of *G_r_* when the function saturates outside the range of possible contrast values. For example, if the best fit function saturates above 100% contrast, maximum value of the function can exceed the largest response seen across the range of contrasts that were actually presented by a substantial margin. Thus, following Itthipuripat et al. (2019), rather than reporting *G_r_* and *G_c_*, we instead obtained a measure of response gain (*R_max_*) by calculating the amplitude of the best-fit Naka-Rushton function at 100% contrast and subtracting the baseline (i.e., *R_max_ = A(100) - b)*, and a measure of contrast gain by calculating the contrast at which the function reaches half the amplitude seen at 100% contrast (*C_50_*).

We used a subject-level resampling procedure to test for differences in the parameters of the fitted Naka-Rushton function across conditions. We drew 100,000 bootstrap samples, each containing *N*-many subjects sampled with replacement, where *N* is the sample size. For each bootstrap sample, we fitted Naka-Rushton function to the amplitude of mean stimulus-evoked CTFs across subjects in the bootstrap sample. We calculated the difference for the parameter between the attend-stimulus and attend-fixation conditions for each bootstrap sample, which yielded a distribution of 100,000 values. We tested whether these difference distributions significantly differed from zero in either direction, by calculating the proportion of values > or < 0, and doubling the smaller value to obtain a 2-sided *p* value.

### Electrode selectivity

We calculated an F-statistic to determine the extent to which responses at each electrode differentiated between spatial positions of the stimulus. For each subject in Experiment 1, we partitioned all data into 15 independent sets (collapsing across the attend-stimulus and attend-fixation conditions, and equating the number of epoch for each stimulus position across sets). We calculated evoked power (averaging across 100-ms windows) for each stimulus position within each set. For each electrode, we calculated the ANOVA F-statistic on evoked power across the eight stimulus positions, with each of the 15 sets serving as an independent observation. Higher F-statistic values indicate that evoked power varied with stimulus position to a greater degree. As with our IEM analyses, we randomly partitioned the data into sets 500 times, and averaged the F-statistic across iterations.

### Data/software availability

All data and code will be made available on Open Science Framework when the manuscript is accepted for publication.

## Results

### Experiment 1

In Experiment 1, we tested how spatial attention modulated spatially selective stimulus-evoked activity measured with EEG. On each trial, observers viewed a series of bullseye stimuli, and we manipulated whether spatial attention was directed toward or away from these stimuli (Fig. 2a). Each trial began with a peripheral cue that indicated where the bullseye stimuli would appear. In *attend-stimulus* blocks, observers covertly monitored the sequence of bullseyes for one bullseye that was lower contrast than the rest. In *attend-fixation* blocks, observers ignored the bullseye stimuli, and instead monitored the fixation dot for a brief decrement in contrast. At the end of each trial, observers reported whether or not a contrast decrement occurred in the attended stimulus. We matched difficulty across the two conditions by adjusting the size of the contrast decrement for each condition (see Materials and Methods, Staircase procedures). Thus, accuracy was comparable in the attend-stimulus (*M* = 81.0%, *SD* = 3.7) and the attend-fixation (*M* = 80.0%, *SD* = 2.2) conditions.

To test how spatial attention modulates the spatial selectivity of stimulus-driven activity, we measured the power of broadband EEG activity evoked by the bullseye stimuli (i.e., the power of activity phase-locked to stimulus onset; see Materials and Methods, Evoked power) and we used an IEM (Brouwer and Heeger, 2009, 2011; Sprague and Serences, 2013; Foster et al., 2016) to reconstruct spatially selective channel-tuning functions (CTFs) from the scalp distribution of stimulus-evoked power (see Materials and Methods, Inverted encoding model). Figure 4a shows stimulus-evoked CTFs across time in the attend-stimulus and attend-fixation conditions. We found that stimulus-evoked CTFs were tuned for the stimulus location, with a peak response in the channel tuned for the stimulus location, and this spatial tuning emerged 70-80 ms after stimulus onset. Human event-related potential (ERP) studies have found that visually evoked responses are modulated by attention as early as 80 ms after stimulus onset (for review, see Hillyard and Anllo-Vento, 1998). For instance, many studies have reported that attention increases the amplitude of the posterior P_1_ component (e.g. van Voorhis and Hillyard, 1977; Martínez et al., 1999; Itthipuripat et al., 2014a), which is typically seen approximately 100 ms after stimulus onset. Thus, we focused our analysis in an early window, 80-130 ms after stimulus onset, to capture the early stimulus-evoked response. Figure 4b shows the reconstructed channel responses during our window of interest for each of the eight stimulus positions, separately for the attend-stimulus and attend-fixation conditions. We found that the peak response was always occurred in the channel tuned for the spatial position of the stimulus. Thus, stimulus position is precisely encoded by stimulus-evoked power. To determine which electrodes carry information about the spatial position of the stimulus, we calculated an F-statistic across stimulus locations for each electrode (see Materials and Methods, Electrode selectivity), where larger values indicate that stimulus-evoked power varies with stimulus location to a greater extent (Fig. 4c). We found that posterior electrodes carried the most information about stimulus location. Although the cortical source of EEG signals cannot be fully resolved based on EEG scalp recordings, this analysis as well as the timing of the observed activity suggest that the spatially selective activity that our IEM analysis capitalized on is generated in posterior visual areas.

**Figure 4.**
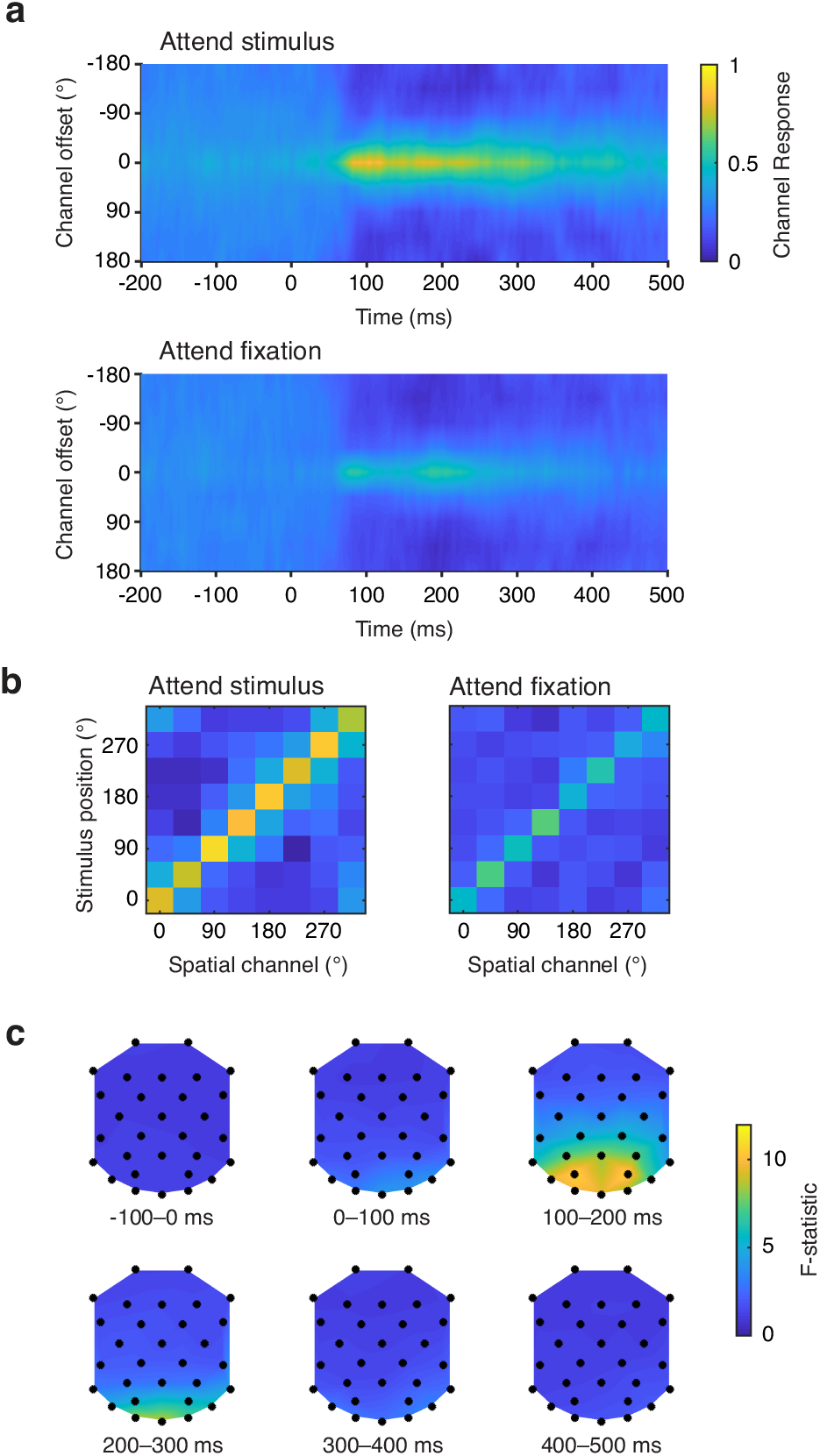
Stimulus-evoked EEG activity encodes stimulus position. **(a)** Time-resolved CTFs reconstructed from stimulus-evoked EEG activity in the attend-stimulus (upper) and attend-fixation (lower) conditions (the stimulus onset at 0 ms). **(b)** Channel responses in our window of interest (80-130 ms after stimulus onset) for each of the eight stimulus positions for the attend-stimulus (left) and attend-fixation (right) conditions. **(c)** Scalp topography of F-statistic values in 100-ms windows (anterior sites are at the top of each topographic plot). Larger values indicate that stimulus-evoked power varies to a greater extent with stimulus position.

Having established that stimulus-evoked power precisely encodes stimulus position, we examined the effect of attention on the tuning properties of the stimulus-evoked CTFs. Figure 5a shows the stimulus-evoked CTFs in our window of interest. We fitted the CTFs in each condition with an exponentiated cosine function to estimate baseline, amplitude, and width parameters (Fig. 2e; Materials and Methods, Model fitting). Figure 5b shows the parameter of the best fitting functions by condition. We found that stimulus-evoked CTFs were both higher in amplitude (*p* < .0001) and more broadly tuned (*p* < .0001) in the attend-stimulus condition than in the attend-fixation condition, and we observed no difference in baseline between the conditions (*p* = .974). However, as we will see next, the finding that CTFs were broader in the attend-stimulus condition than in the attend-fixation condition appears to be an artifact of lingering activity from the preceding stimulus event. Furthermore, this effect did not replicate in Experiment 2. Thus, the primary effect of attention is to improve the stimulus representation via an increase in the amplitude of the CTF that tracks the target’s position.

#### Controlling for lingering activity evoked by the preceding stimulus in the sequence

We designed our task to measure activity evoked by each of the four stimuli presented within each trial. To this end, we jittered the inter-stimulus interval between each stimulus (between 500 and 800 ms) to ensure that activity evoked by one stimulus in the sequence will not be phase-locked to the onsets of the stimuli before or after it in the sequence. However, when we examined the amplitude of stimulus-evoked CTFs through time (Fig. 5c), we found pre-stimulus tuning (in the 200 ms preceding stimulus onset) that was higher amplitude in the attend-stimulus than attend-fixation condition (*p* = .036). We hypothesized that this pre-stimulus spatially selective activity may reflect activity evoked by the preceding stimulus in the sequence that was sufficiently low frequency that was not eliminated by the temporal jitter between stimulus onsets. Because this pre-stimulus activity was higher amplitude in the attend-stimulus condition than in the attend-fixation condition, it could have contaminated the apparent attentional modulations of stimulus-evoked activity (both the increase in amplitude and the broadening of stimulus-evoked CTFs) that we observed 80-130 ms after stimulus onset. Thus, we examined the effect of this lingering activity by examining CTFs as a function of position in the sequence of four stimuli within each trial. Within each trial, the second, third, and fourth stimuli were preceded by a bullseye stimulus that should drive a strong visually evoked response, whereas the first stimulus was preceded by a small, low-contrast cue that should drive a much weaker visually evoked response (see Fig. 1). Thus, we expected that stimulus-evoked activity for the first bullseye stimulus in the sequence should be contaminated by activity evoked by the preceding stimulus to a lesser degree than subsequent stimuli in the sequence. Figure 6 shows the reconstructed CTFs from activity evoked by stimuli in each position on the sequence. For this analysis, we trained the IEM on all but the tested stimulus. For example, when testing on the first stimulus in the sequence, we trained on stimuli in serial positions 2-4. We found a robust effect of attention on the amplitude of the stimulus-evoked CTFs across stimuli in all positions in the sequence (all *p*s < .05). In contrast, we found that the CTFs were broader in the attend-stimulus and attend-fixation conditions for the second, third, or fourth stimuli in the sequence (all *p*s < .05), but not for the first stimulus in the sequence (*p* = .540), when the influence of lingering stimulus-evoked activity should be greatly reduced. This finding suggests that the increase in CTF width was driven by lingering activity evoked by the preceding stimulus in the sequence. It is not entirely clear why lingering activity from the preceding stimulus increased the width of CTFs rather than simply increasing CTF amplitude. One possibility is that spatially tuned activity evoked by a visual stimulus is more broadly tuned at later latencies than during the initial encoding of the stimulus.

**Figure 5.**
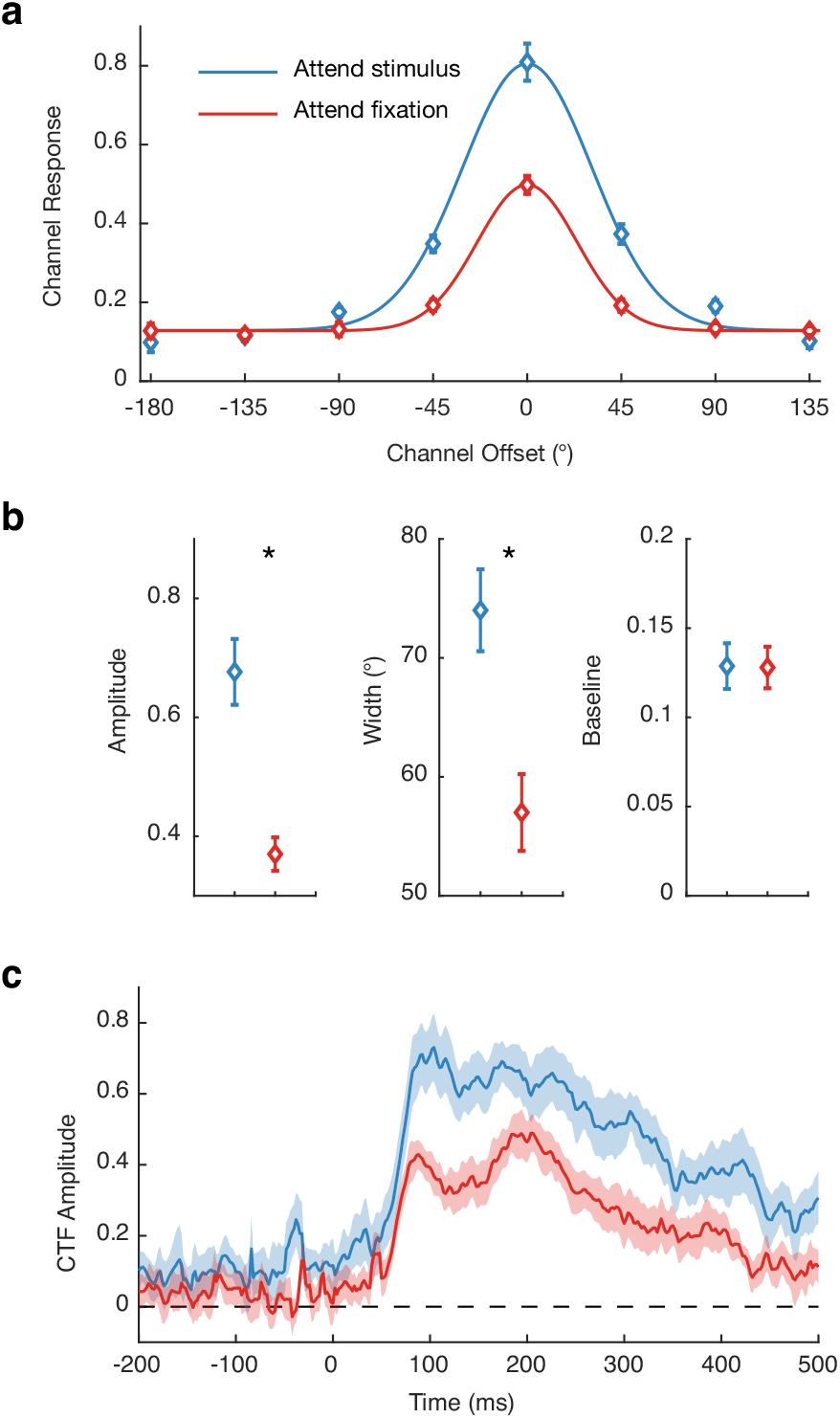
Spatial attention increases the amplitude of stimulus-evoked CTFs. **(a)** Stimulus-evoked CTFs (measured 80-130 ms after stimulus onset) for the attend-stimulus (blue) and attend-fixation (red) conditions. The curves show the best fitting functions. **(b)** Amplitude, width, and baseline parameters of the best fitting functions by for each condition. Asterisks mark differences between the conditions that were significant at the .05 level. **(c)** Amplitude of stimulus-evoked CTFs as a function of time (stimulus onset at 0 ms). All error bars show ±1 bootstrapped SEM.

**Figure 6.**
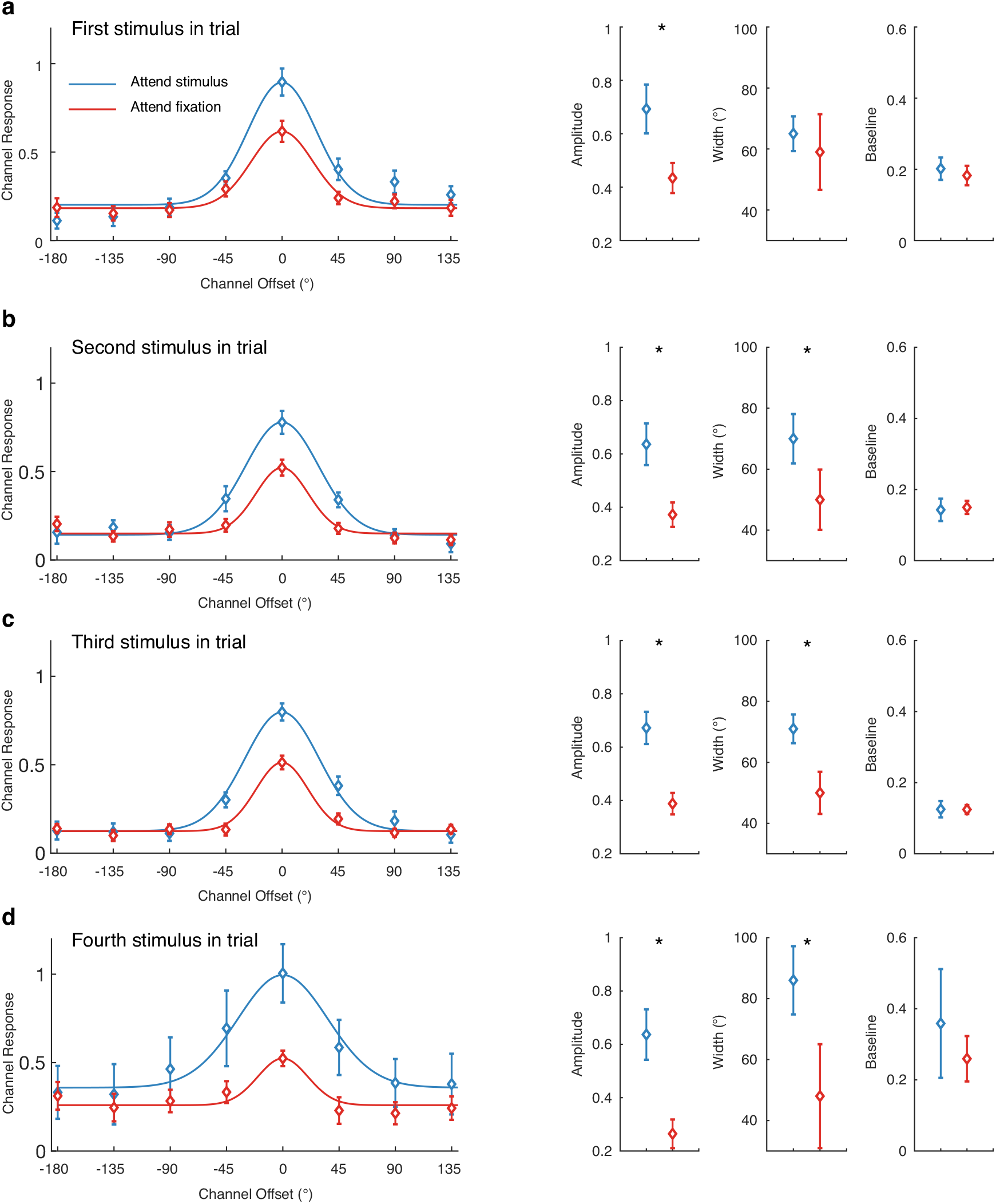
Stimulus-evoked CTFs for each stimulus in the trial sequence. Stimulus-evoked CTFs (measured 80-130 ms after stimulus onset) with best fitting functions (left) and parameter estimates of the best fitting functions (right). Asterisks mark differences between the conditions that were significant at the .05 level. Error bars show ±1 bootstrapped SEM.

Next, to obtain converging evidence for this conclusion, we took a different approach to eliminate lingering activity evoked by the preceding stimulus while still collapsing across all stimulus positions in the sequence. It is primarily low-frequency components that survive temporal jitter. Thus, we reanalyzed the data, this time applying a 4-Hz high-pass filter to remove very low-frequency activity. We found that high-pass filtering the data eliminated the pre-stimulus difference in spatial selectivity between the attend-stimulus and attend-fixation conditions (*p* = .458, see Fig. 7c), suggesting that the pre-stimulus activity was restricted to low frequencies. Having established that a high-pass filter eliminated pre-stimulus activity, we re-examined stimulus-evoked CTFs in our window of interest (80-130 ms) after high-pass filtering (Fig. 7a and 7b). Again, we found that the CTFs were higher amplitude when the stimulus was attended (*p* < .0001). We also found that CTFs were more broadly tuned when the stimulus was attended (*p* < .01). However, as we will see, this small effect of attention on CTF width did not replicate in Experiment 2, suggesting that the primary effect of attention is to increase the amplitude of stimulus-evoked CTFs.

**Figure 7.**
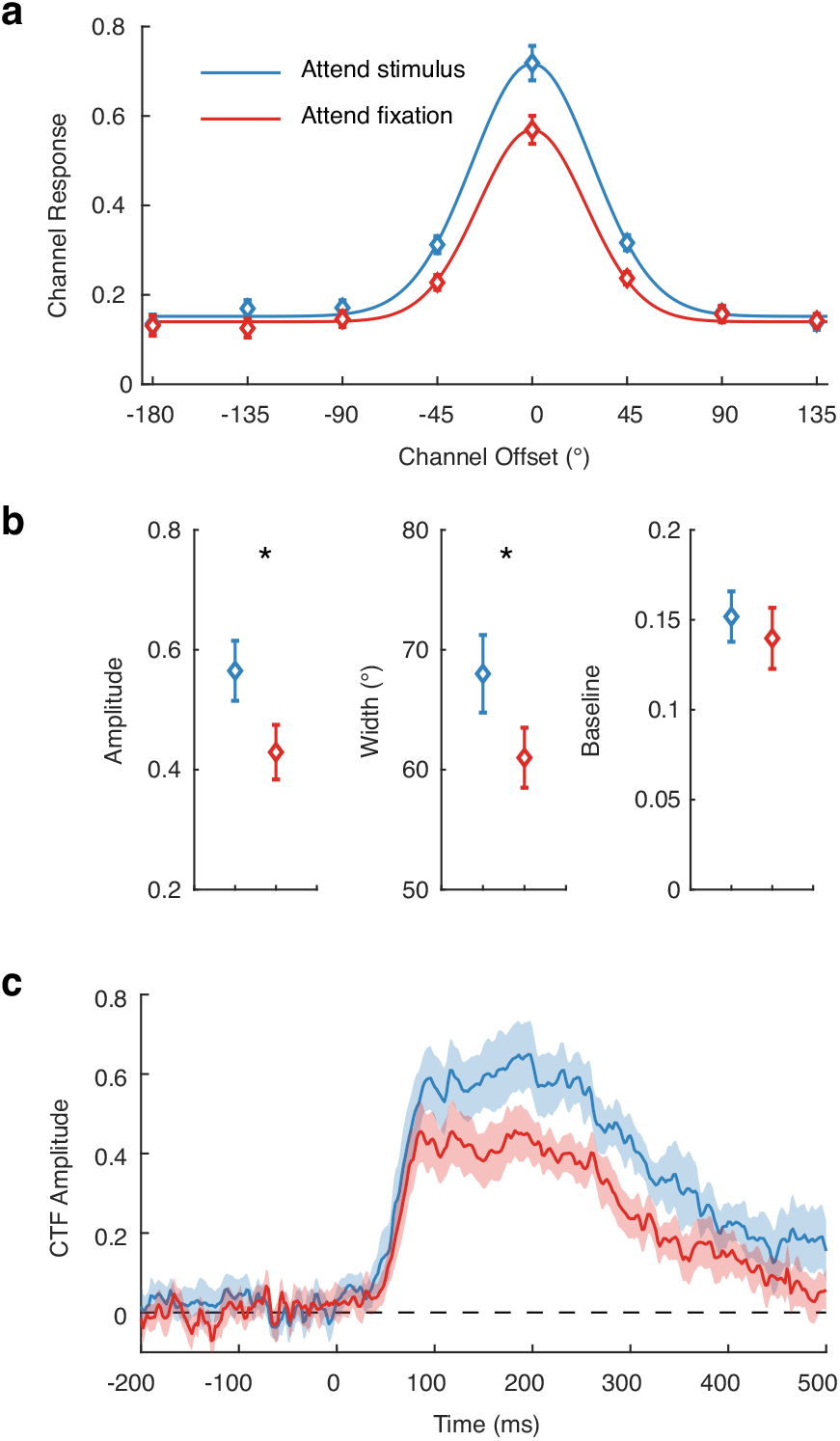
Stimulus-evoked CTFs after high-pass filtering to remove lingering activity from the preceding stimulus. **(a)** Stimulus-evoked CTFs (measured 80-130 ms after stimulus onset) for the attend-stimulus (blue) and attend-fixation (red) conditions. The curves show the best fitting functions. **(b)** Amplitude, width, and baseline parameters of the best fitting functions by for each condition. Asterisks mark differences between the conditions that were significant at the .05 level. **(c)** Amplitude of stimulus-evoked CTFs as a function of time (stimulus onset at 0 ms). All error bars show ±1 bootstrapped SEM.

### Experiment 2

Past fMRI work has found that spatially attending a stimulus increases the amplitude of spatial representations in visual cortex (Sprague and Serences, 2013; Vo et al., 2017). However, this effect of attention on the amplitude of this spatially specific activity is additive with stimulus contrast, such that attention effects are equivalent across all levels of stimulus contrast (Buracas and Boynton, 2007; Murray, 2008; Sprague et al., 2018b; Itthipuripat et al., 2019). Therefore, these changes in spatially specific activity measured with fMRI appear to reflect a stimulus-independent, additive shift in cortical activity that does not provide insight into how attention affects stimulus-evoked sensory processing. In contrast, the CTFs reconstructed from stimulus-evoked EEG activity provides a more direct window into how attention affects *stimulus-driven* sensory activity by isolating activity that is phase-locked to target onset. Therefore, in Experiment 2, we manipulated stimulus contrast to test how the effect of of attention on stimulus-evoked population codes scales with stimulus contrast.

Observers performed the same task as in Experiment 1 (Fig. 2a), but we parametrically varied the pedestal contrast of the bullseye stimulus from 6.25 to 90.6% across trials. We adjusted the size of the contrast decrement independently for each of the conditions using a staircase procedure designed to hold accuracy at approximately 76% correct (see Materials and Methods, Staircase procedures). Accuracy was well matched across condition: mean accuracy across subjects did not deviate from 76% by more than 1% any condition (Table 2). We reconstructed CTFs independently for each condition, having first estimated channel weights using additional trials (with a pedestal contrast of 90.6%) that were collected for this purpose (see Materials and Methods, Training and testing data). In Experiment 2, we again used a 4-Hz high-pass filter to remove lingering activity evoked by the preceding stimulus in the sequence. Figure 8a and 8b show the stimulus-evoked CTFs as a function of contrast with the best-fit functions for the attend-stimulus and attention-fixation conditions, respectively. For each of the three parameters (amplitude, baseline, and width) we performed a resampling test to test for a main effect of contrast, a main effect of attention, and an attention × contrast interaction (see Materials and Methods, Resampling tests). First, we examined CTF amplitude (Fig. 8c). We found that CTF amplitude increased with stimulus contrast (main effect of contrast: *p* < .0001), and CTF amplitude was larger in the attend-stimulus condition than in the attend-fixation condition (main effect of attention: *p* < .0001). Critically, the effect of attention on CTF amplitude increased with stimulus contrast (attention × contrast interaction, *p* < .0001). This finding provides clear evidence that the effect of attention on stimulus-evoked CTFs is not additive with stimulus contrast, as is the case with BOLD activity measured by fMRI (Buracas and Boynton, 2007; Murray, 2008; Sprague et al., 2018b; Itthipuripat et al., 2019).

**Table 2.** Mean accuracy (and standard deviation) in Experiment 2 as a function of task condition and pedestal contrast of the bullseye stimuli.

| Pedestal contrast | 6.25% | 12.5% | 25.0% | 50.0% | 90.6% |
| --- | --- | --- | --- | --- | --- |
| Attend stimulus | 75.4% (0.97) | 75.8% (0.80) | 76.1% (0.96) | 75.5% (0.60) | 76.4% (0.50) |
| Attend fixation | 76.0% (0.86) | 76.0% (0.97) | 76.0% (0.74) | 76.1% (0.85) | 76.1% (0.31) |

**Figure 8.**
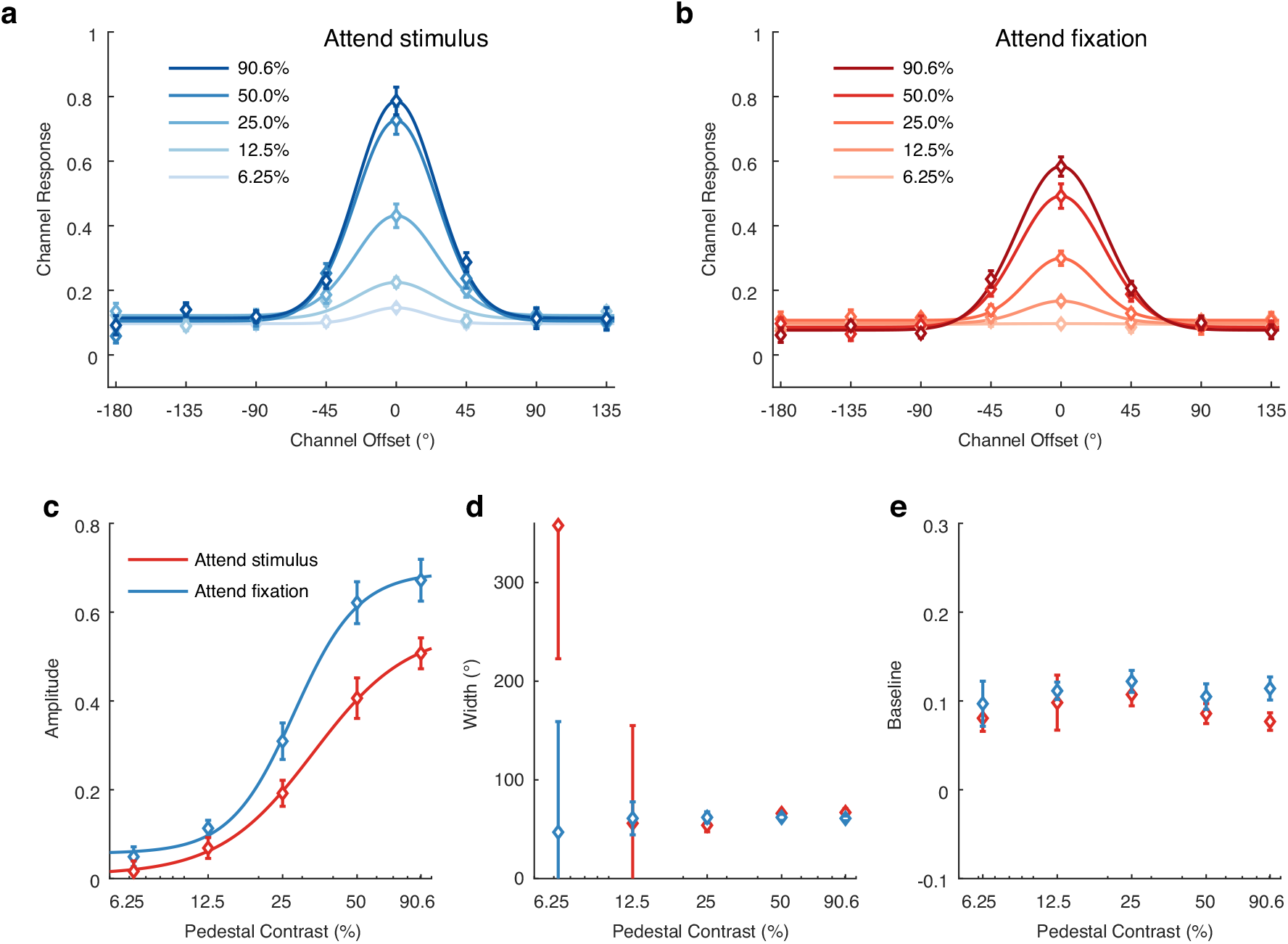
The effect of spatial attention on the amplitude of stimulus-evoked CTFs scales with stimulus contrast. **(a-b)** Stimulus-evoked CTFs (measured 80-130 ms after stimulus onset) as a function of stimulus contrast in the attend-stimulus and attend-fixation conditions in Experiment 2. Curves show the best-fit exponentiated cosine functions. **(c-e)** Amplitude, width (fwhm), and baseline parameters of stimulus-evoked CTFs as a function of task condition and stimulus contrast. Curves in **(c)** show the best-fit Naka-Rushton function to CTF amplitude. Error bars reflect ±1 bootstrapped SEM across subjects.

To further characterize this effect, we fitted the amplitude parameter with a Naka-Rushton function (Materials and Methods, Quantifying contrast-response functions). The curves in Figure 8c show the best-fit functions for each condition. We estimated four parameters of the Naka-Rushton function: a baseline parameter (*b*), which determines the offset of the function from zero, a response gain parameter (*R_max_)*, which determines how much the function rises above baseline, and contrast gain parameter (*C_50_*), which measures horizontal shifts in the function, and a slope parameter (*n)*, which determines how steeply the function rises. We found that *R_max_* was reliably higher in the attend-stimulus condition the attend-fixation condition (resampling test, *p* = 0.036). However, we did not find reliable differences between conditions for the *C_50_, b*, or *n* parameters (resampling tests, *p* = 0.104, *p* = 0.126, *p* = 0.376, respectively, see Table 3 for descriptive statistics). Thus, we found that attention primarily changed the amplitude of stimulus-evoked CTFs via an increase in response gain.

**Table 3.** Mean (and bootstrapped *SEM)* of the parameter estimates from the Naka-Rushton fits to the amplitude of stimulus-evoked CTFs in Experiment 2.

| Parameter | $R_{max}$ | $C_{50}$ | $b$ | $n$ |
| --- | --- | --- | --- | --- |
| Attend stimulus | 0.62 (0.05) | 26.18 (1.27) | 0.06 (0.02) | 3.35 (1.66) |
| Attend fixation | 0.51 (0.04) | 30.79 (3.73) | 0.01 (0.02) | 2.27 (1.23) |

Next, we examined CTF width (Fig. 8d). We found that estimates of CTF width were very noisy for the 6.25% and 12.5% contrast conditions because of the low amplitude of the CTFs in these conditions, precluding confidence in those estimates. Thus, we restricted our analysis to the higher contrast conditions (25.0, 50.0, and 90.6% contrast). We found no main effect of attention (*p* = .851), and no main effect of contrast (*p* = .130). However, we found a reliable attention × contrast interaction (*p* = .035), such that CTFs were narrower when the stimulus was attended for the 90.6% contrast condition and 50% contrast condition, and were broader for the 25% contrast condition, but none of these differences between the attend-stimulus and attend fixation conditions survived Bonferoni correction (*p* = .043, *p* = .277, and *p* = .258, respectively; *α*_corrected_ = .05/3 = .017). Thus, we did not replicate the finding from Experiment 1 that stimulus-evoked CTFs were broader when the stimulus was attended. Finally, we examined CTF baseline (Fig. 8e). Although CTF baseline was generally higher in the attend-stimulus condition than in this attention fixation condition, this difference was not significant (main effect of attention, *p* = .055), nor was the main effect of contrast (*p* = .708) or attention × contrast interaction (*p* = .289)

### Attention produces a baseline shift in spatially selective alpha-band power

Past work has closely linked alpha-band (8–12 Hz) oscillations with covert spatial attention. A plethora of studies has shown that posterior alpha-band power is reduced contralateral to an attended location (e.g. Worden et al., 2000; Kelly et al., 2006; Thut et al., 2006). Furthermore, alpha-band activity precisely tracks where in the visual field spatial attention is deployed (Rihs et al., 2007; Samaha et al., 2016; Foster et al., 2017). For example, we and others have reconstructed spatial CTFs from alpha-band activity that track the spatial and temporal dynamics of covert attention (e.g. Foster et al., 2017). Importantly, the relationship between alpha topography and attention appears to include a stimulus-independent component, because alpha activity tracks the allocation of spatial attention in blank or visually balanced displays (Worden et al., 2000; Thut et al., 2006). More recent work has provided further evidence in favor of this view. Itthipuripat et al. (2019) parametrically varied the contrast of a lateral stimulus and cued observers to either attend the stimulus or attend the fixation dot (similar to the task we use in the current study). Itthipuripat and colleagues found that the effect of attention and stimulus contrast on posterior alpha-band power contralateral to the stimulus were additive: although contralateral alpha power declined as stimulus contrast increased, directing attention to the stimulus reduced contralateral alpha power by the same margin regardless of stimulus contrast. This finding suggests that the alpha-band activity indexes the locus of spatial attention in a stimulus-independent manner.

If alpha-band activity reflects a stimulus-independent aspect of spatial attention, then fluctuations of alpha power should be additive with stimulus contrast in Experiment 2. Thus, we examined CTFs reconstructed from total alpha-band power (i.e. the power of alpha-band activity regardless of its phase relationship to stimulus onset) in a post-stimulus window (0-500 ms after stimulus-onset). Figure 9a and 9b show the reconstructed alpha-band CTFs for the attend-stimulus and attend-fixation conditions, respectively. Figures 9c-e show the amplitude, width, and baseline parameters as a function of condition. We found that amplitude of alpha-band CTFs (Fig. 9c) increased with stimulus contrast (main effect of contrast: *p* < .0001), and CTF amplitude was greater in the attend-stimulus condition than in the attend-fixation condition (main effect of attention: *p* = 0.0005). Importantly, we did not find a reliable interaction between attention and stimulus contrast on CTF amplitude (attention × contrast interaction, *p* = 0.438). Thus, the effects of contrast and attention on the amplitude of alpha CTFs was additive. Although spatial CTFs were generally broader in the attend-stimulus condition than in the attend-fixation condition (Fig. 9d), we did not find a reliable main effect of attention (*p* = 0.094), nor did we find a main effect of contrast (*p* = 0.869) or an attention x contrast interaction (*p* = 0.908). Finally, we found that baseline was reliably lower in the attend-stimulus condition than in the attend-fixation condition (Fig. 9e, main effect of attention: *p* < .001). Thus, attending the stimulus not only increased activity in the channel tuned for the attended location, but also reduced activity in channels tuned for distant locations. We did not find a reliable main effect of contrast (p = 0.080), or an attention x contrast interaction (*p* = 0.900). To summarize, spatial attention primarily influenced the amplitude and baseline of alpha-band CTFs, and these effects were additive with the effect of stimulus contrast. Thus, the effect of attention of alpha-band power reflects a stimulus-independent baseline shift in spatially selective alpha-band power, much like the effect of attention on spatially-specific BOLD activity in past fMRI studies of attention (Murray, 2008; Itthipuripat et al., 2019).

**Figure 9.**
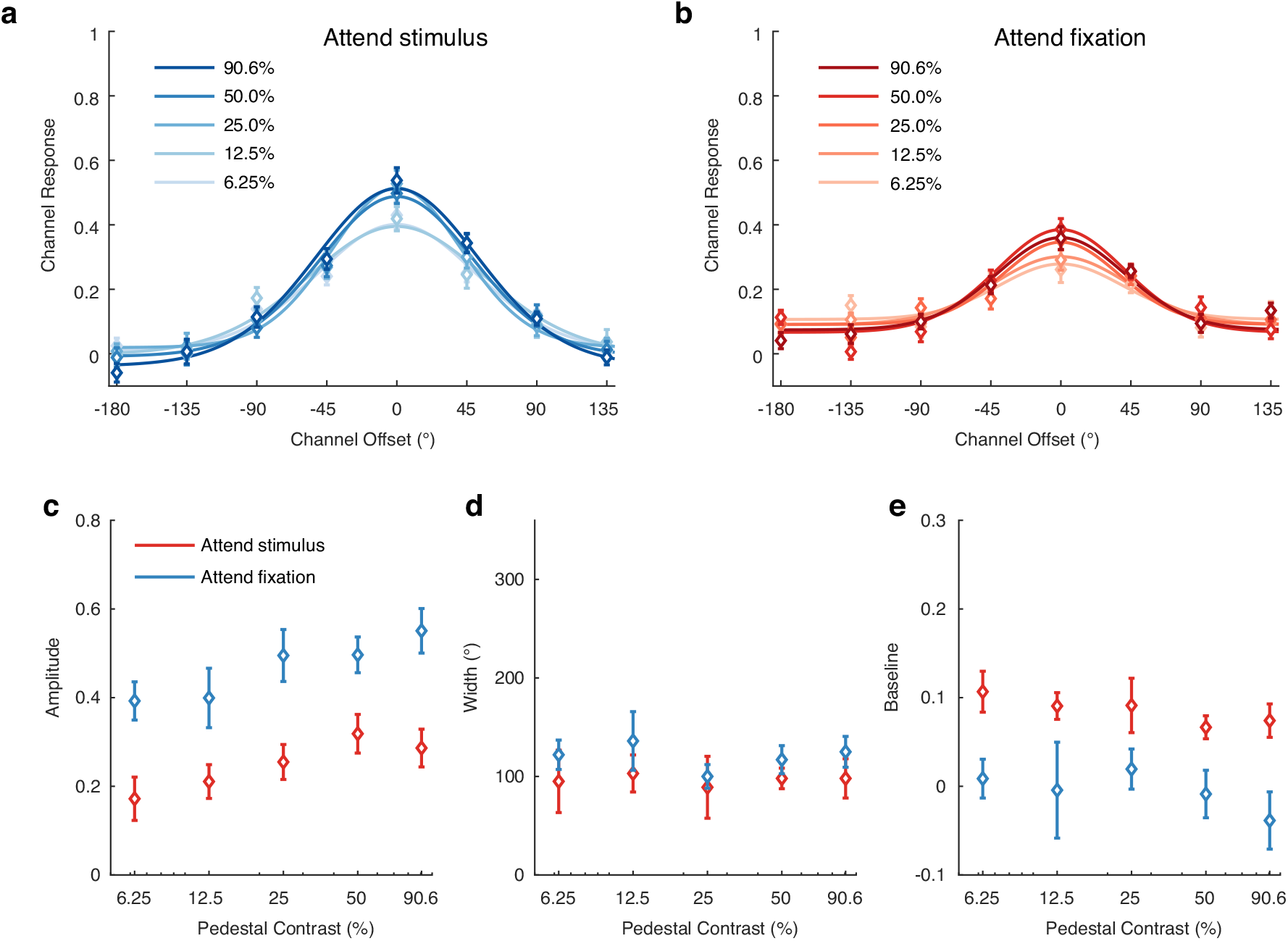
Spatial attention produces an additive shift in the amplitude of alpha-band CTFs. **(a-b)** Alpha-band CTFs (measured 0-500 ms after stimulus onset) as a function of stimulus contrast in the attend-stimulus and attend-fixation conditions in Experiment 2. Curves show the best-fit exponentiated cosine functions. **(c-e)** Amplitude, width (fwhm), and baseline parameters of alpha-band CTFs as a function of task condition and stimulus contrast. Error bars reflect ±1 bootstrapped SEM across subjects.

## Discussion

To examine how and when covert spatial attention shapes the selectivity of stimulus-driven spatial population codes, we reconstructed spatially selective channel tuning functions from stimulus-evoked EEG signals that were phase-locked to stimulus onset. Across two experiments, we found that attention increased the amplitude of stimulus-evoked CTFs that were tuned for the location of the stimulus. We did not find convincing evidence that attention changed the width of stimulus-evoked CTFs. Although we found that stimulus-evoked CTFs were broader for attended stimuli than for unattended stimuli in Experiment 1, this effect was greatly reduced when the influence of prior stimulus events was accounted for, and did not replicate in Experiment 2. Therefore, our results show that spatial attention primarily increases the amplitude of stimulus-evoked population tuning functions.

A core strength of our EEG-based approach is that it allowed us to isolate early visually evoked activity. We focused our analysis on stimulus-evoked activity in a window 80-130 ms after stimulus onset. Visually evoked EEG activity at this latency reflects the first wave of stimulus-driven activity in extrastriate cortex (Clark and Hillyard, 1996; Martínez et al., 1999), but likely also captures early recurrent feedback signals (e.g. Boehler et al., 2008). Many ERP studies have shown that spatial attention increases the amplitude of evoked responses at this early latency. For example, spatial attention increases the amplitude of the posterior P1 component observed approximately 100 ms after stimulus onset (van Voorhis and Hillyard, 1977; Martínez et al., 1999; Itthipuripat et al., 2014a). However, it is unclear how changes in the overall amplitude of visually evoked potentials correspond to changes in underlying population codes. For instance, a larger overall population response could reflect an increase in the amplitude of the spatial population code, or it could reflect a broadening of the spatially tuned population response without increasing its amplitude, such that the stimulus evoked a response in a larger population of neurons. Here, we provide the first clear evidence that attention enhances the amplitude of the stimulus-evoked spatial population codes during this early stage of sensory processing.

In Experiment 2, we confirmed that we were observing an attentional modulation of stimulus-evoked activity rather than a stimulus-independent increase in baseline activity. Here, we found that the effect of attention on the amplitude of stimulus-evoked CTFs increased with stimulus contrast. Model fitting revealed that this effect was best described by an increase in response gain (i.e., a multiplicative scaling of the CRF), which dovetails with past work that has found that attention increases response gain of the P1 component and of steady-state visually evoked potentials (Kim et al., 2007; Itthipuripat et al., 2014a, 2014b, 2019). Although our results are most consistent with an increase in response gain, it must be noted that our CRFs did not clearly saturate at higher stimulus contrast, which makes it difficult to unambiguously differentiate between response gain and contrast gain because contrast gain can mimic response gain in the absence of clear saturation (e.g. consider the left half of the functions in Fig. 1b, which closely resemble a change in response gain). We also note that our finding that attention increased response gain may depend on the fact that we cued the precise location of the bullseye stimulus. The normalization model of attention (Reynolds and Heeger, 2009), an influential computational model of attention, predicts that whether attention produces a change in response gain or contrast gain depends on the spread of spatial attention relative to the size of the stimulus. Specifically, the model predicts that attention will change response gain when attention is tightly focused on a stimulus, but will change contrast gain (shifting the CRF to the left) when the spatial spread of attention is large relative to the stimulus (Reynolds and Heeger, 2009). Indeed, past EEG and psychophysical studies that have manipulated the spatial spread of attention relative to the size of the stimulus have supported this prediction (Herrmann et al., 2011; Itthipuripat et al., 2014b). Thus, further work is needed to test whether the change in response gain that we observed in the amplitude of the spatially tuned population response is specific to situations in which observers can focus spatial attention very tightly on the stimulus. Nevertheless, Experiment 2 provides unambiguous evidence that the effect of attention on the amplitude of spatially tuned population responses reflects a modulation of stimulus-driven activity rather than a stimulus-independent, additive shift as is measured with fMRI (Buracas and Boynton, 2007; Murray, 2008; Pestilli et al., 2011; Sprague et al., 2018b; Itthipuripat et al., 2019; but see Li et al., 2008).

Other aspects of our findings, however, are consistent with the stimulus-independent effects that have been observed in BOLD activity. There is substantial evidence that attention is linked with spatially specific changes in alpha-band power (for reviews, see Jensen and Mazaheri, 2010; Foster and Awh, 2019). Many studies have shown that alpha power is reduced contralateral to attended locations (e.g. Worden et al., 2000; Thut et al., 2006). This reduction is thought to reflect a stimulus-independent shift in alpha power because it is seen in in the absence of visual input (Sauseng et al., 2005; Foster et al., 2020). Recently, Itthipuripat et al. (2019) provided new support for this view. They found that spatially attending a lateralized stimulus reduced alpha power by the same margin regardless of stimulus contrast. We conceptually replicated and extended this finding. Attention related modulations of alpha power track the precise location that is attended within the visual field (Rihs et al., 2007; Samaha et al., 2016; Foster et al., 2017). Thus, we examined the effect of attention on post-stimulus alpha-band CTFs. Consistent with Itthipuripat et al.’s (2019) results, we found that the effect of attention on post-stimulus alpha-band CTFs was additive with the effect of stimulus contrast, such that spatial attention increased the amplitude of spatially tuned alpha-band CTFs by the same amount regardless of stimulus contrast. Thus, our results add to growing evidence that attention-related changes in alpha-band power are stimulus independent.

## Conclusions

Decades of work have established that spatial attention modulates relatively early stages of sensory processing, but there has been limited evidence regarding how attention changes population-level sensory codes. Here, we have provided robust evidence that spatial attention increases the amplitude of spatially-tuned neural activity evoked by attended items within 100 ms of stimulus onset. Thus, attention increases the gain of spatial population codes during the first wave of sensory activity.

## Conflicts of interest

None

## Author contributions

JJF and EA conceived of the experiments. JJF designed the experiments. JJF, WT, and JWW carried out the experiments. JJF analyzed the data (including writing code for data analysis). WT vetted analysis code. JJF drafted the manuscript. All authors revised the manuscript, and approved the final version for submission.

## Acknowledgements

This work was supported by National Institute of Mental Health Grant 5RO1 MH087214-08. We thank Mei Arditi, Emma Bsales, and Naomi Nero for assistance with data collection.

## Notes

### Competing Interest Statement

The authors have declared no competing interest.

